# Protection against reinfection with D614- or G614-SARS-CoV-2 isolates in hamsters

**DOI:** 10.1101/2021.01.07.425729

**Authors:** Marco Brustolin, Jordi Rodon, María Luisa Rodríguez de la Concepción, Carlos Ávila-Nieto, Guillermo Cantero, Mónica Pérez, Nigeer Te, Marc Noguera-Julián, Víctor Guallar, Alfonso Valencia, Núria Roca, Nuria Izquierdo-Useros, Julià Blanco, Bonaventura Clotet, Albert Bensaid, Jorge Carrillo, Júlia Vergara-Alert, Joaquim Segalés

## Abstract

Reinfections with SARS-CoV-2 have already been documented in humans, although its real incidence is currently unknown. Besides having great impact on public health, this phenomenon raises the question if immunity generated by a single infection is sufficient to provide sterilizing/protective immunity to a subsequent SARS-CoV-2 re-exposure. The Golden Syrian hamster is a manageable animal model to explore immunological mechanisms able to counteract COVID-19, as it recapitulates pathological aspects of mild to moderately affected patients. Here, we report that SARS-CoV-2-inoculated hamsters resolve infection in the upper and lower respiratory tracts within seven days upon inoculation with the Cat01 (G614) SARS-CoV-2 isolate. Three weeks after primary challenge, and despite high titers of neutralizing antibodies, half of the animals were susceptible to reinfection by both identical (Cat01, G614) and variant (WA/1, D614) SARS-CoV-2 isolates. However, upon re-inoculation, only nasal tissues were transiently infected with much lower viral replication than those observed after the first inoculation. These data indicate that a primary SARS-CoV-2 infection is not sufficient to elicit a sterilizing immunity in hamster models but protects against lung disease.

## Introduction

The Severe Acute Respiratory Syndrome Coronavirus 2 (SARS-CoV-2) is the etiological agent of Coronavirus Infectious Disease 2019 (COVID-19), a respiratory affection that spread globally with an unprecedented rapidity and severity, impacting both international public health and economics. To date, SARS-CoV-2 has infected more than 84 million people globally, resulting in more than 1.8 million deaths, as reported by the World Health Organization ^1^.

Unraveling immunopathological disorders caused by SARS-CoV-2 is one of the priorities of the scientific community. SARS-CoV-2 infection induces a rapid production of neutralizing antibodies ^2^; however, the magnitude of a neutralizing response as well as its decay correlates directly with the severity of the disease. To date, longitudinal studies confirmed the duration of neutralizing antibodies for 2 to 5 months post-symptoms onset ^3–6^. In addition, the degree of protection against a reinfection event caused by identical or other viral variants is still not clear. The first evidence of COVID-19 reinfection was described by August 25^th^, 2020 ^7^. This study reported reinfection of an individual 142 days after the first infectious episode. The patient did not display any symptom during the second infection; viruses belonging to different SARS-CoV-2 clades were identified from the first and second episodes. Immediately after this first case, other SARS-CoV-2 reinfections have been reported in several countries, including The Netherlands, Belgium, Spain, Sweden, Qatar, South Korea, United States, Ecuador and India ^8–12^. The symptoms described in these cases had different degrees of severity compared to the first infectious event and ranged from asymptomatic to severe disease, being more intense during the second infections in few patients. In all cases, differences in viral genomic sequences were identified between the first and second infections.

Experimental reinfection studies have been performed in non-human primates (NHPs), transgenic mice expressing the human angiotensin-converting enzyme 2 (hACE2), Cyclophosphamide (CyP) immunosuppressed and RAG2-knockout Golden Syrian hamster (*Mesocricetus auratus*) and cat (*Felis catus*) ^13–17^. In all models, animals re-challenged with the same SARS-CoV-2 isolate developed a protective non-sterilizing immunity. Currently, there is no data available about the induction of protective immunity conferred by a given strain *versus* another variant.

The Golden Syrian Hamster is a suitable model to study COVID-19 ^18,19^. SARS-CoV-2 can replicate on both upper and lower respiratory tracts in this animal model. Upon challenge, animals develop a mild-to-moderate disease with a recovery period ranging from one to two weeks. Importantly, infection with SARS-CoV-2 in hamsters recapitulates several lesions observed in the human lower respiratory tract. These include pneumonia with bilateral lungs involvement, ground-glass opacities, presence of focal oedema, inflammation, and acute respiratory distress syndrome ^20^. To date, excluding hamsters, only NHPs partially reproduce the clinical picture experienced by COVID-19 human patients. In addition, age and sex-linked differences in SARS-CoV-2 infection and clinical signs have been reported in hamsters, reflecting human similarities ^18,21^. Thus, the golden Syrian hamster could be an appropriate model to study SARS-CoV-2 reinfections.

Here, we test the capacity of SARS-CoV-2 to reinfect golden Syrian hamsters using two variants of the virus: Cat01, a variant isolated from a human patient in Spain and WA/1 a variant isolated from a human patient in the USA. The Cat01 isolate differs from the WA/1 one by the presence of 15 single point mutations. Among them, the most striking is at the 614 position of the Spike protein gene; the WA/1 isolate possess a wild-type D614 spike protein, while the Cat01 isolate displays the G614 mutation. S-G614 strains emerged for the first time in Europe during March 2020 and quickly spread globally, arriving almost at fixation and replacing S-D614 variants ^22^. Further studies demonstrated that D614G variants have a higher transmission capacity ^23–25^ and reach higher viral loads in the upper airways ^26^. It is therefore important to gain insights into mechanisms of reinfection and the development of protective immunity using different viral strain, which could interfere with a primary infection event.

Our results demonstrate that animals exposed to Cat01 variant developed a cross-protective but not sterilizing immune response against a second infection event, regardless of the viral variant used for the re-challenge. Importantly, we showed that identical and variant viral strains could successfully infect the upper respiratory tract of re-challenged animals, but no evidence of infection occurred at the lower respiratory tract.

## Results

### Hamsters do not lose weight after SARS-CoV-2 reinfection

The level of protection conferred by prior inoculation of SARS-CoV-2 was explored. Weight loss was used to clinically track infection and reinfection ^15^ (Figure 1). Hamsters inoculated for the first time with 10^5.8^ TCID_50_/animal of SARS-CoV-2 isolate Cat01 (n=24) showed a progressive reduction of weight starting from 1 until 5 day post infection (dpi) compared to the mock-inoculated animals (Figure 1A), according to what has been previously described ^13,19^. The maximum mean weight loss for infected animals was - 3.56% (SD ± 4.34%; *n* = 16) at 5 dpi. From 6 dpi onwards, animals recovered weight in a similar trend to that of the control group, indicating the beginning of the clinical recovery. Mean weight differences between SARS-CoV-2 and mock-infected animals were statistically significant starting from 2 dpi. On day 21 after inoculation, twelve animals that underwent a primary SARS-CoV-2 infection with the Cat01 isolate were re-challenged with Cat01 (n= 6) or WA/1 (n= 6) isolates, at a final concentration of 10^5.2^ TCID_50_/animal. Upon reinfection with both variants, animals did not have statistically significant weight variations compared to the control group (Figure 1B).

**Figure 1.**
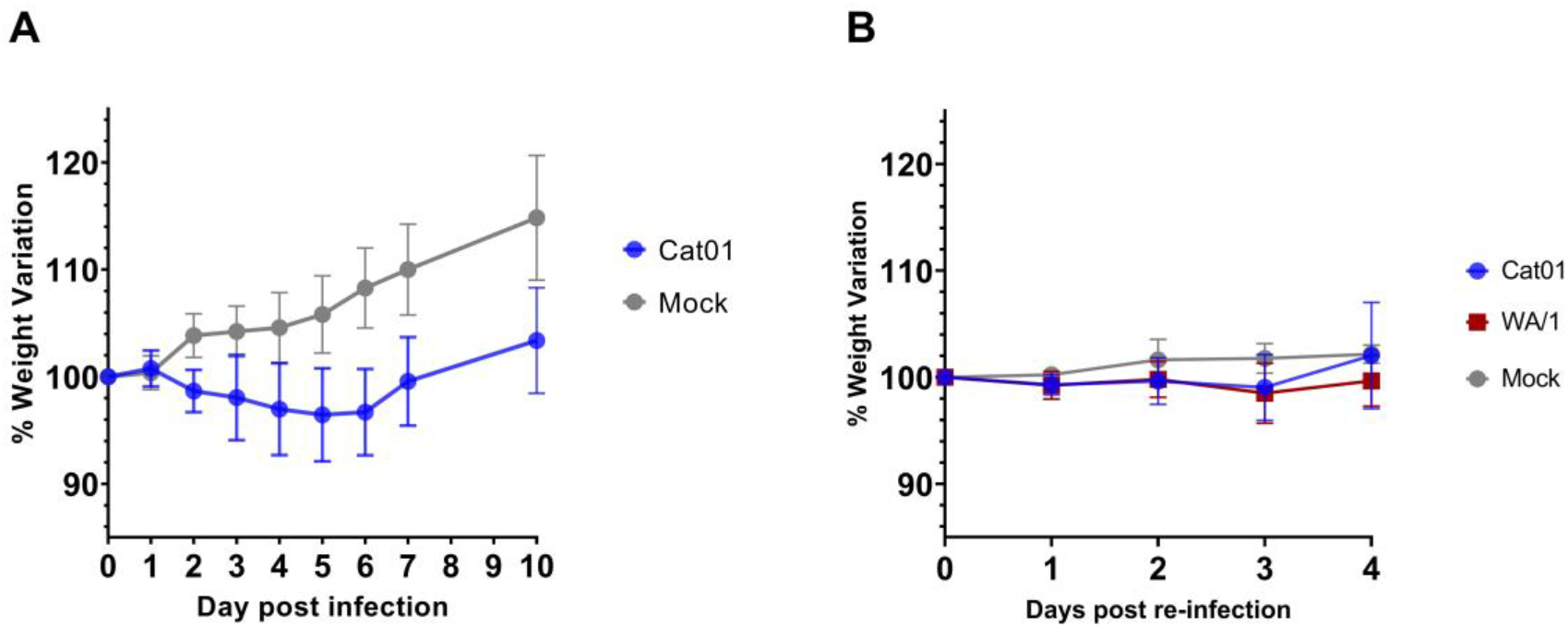
Weight variation upon first inoculation and re-challenge. Data are expressed as percentage of variation referred to the weight recorded at the day of the challenge (a) or re-challenge (b). **a)** Mean percentage of weight variation of animals inoculated with SARS-CoV-2 Cat01 variant (blue) or with PBS mock solution (grey). **b)** Mean percentage of weight variation of animals after SARS-CoV-2 re-inoculation. In blue animals exposed to SARS-CoV-2 Cat01 variant, in red animals exposed to SARS-CoV-2 WA/1 variant, and in grey animals exposed to PBS mock solution.

### SARS-CoV-2 causes rhinitis and bronchointersticial pneumonia upon infection but minimal to no lesions upon reinfection

Moderate to severe inflammatory lesions were observed in nasal turbinates at 2 and 4 dpi, being mild at 7 dpi. Animals developed multifocal to diffuse, muco-purulent to non-suppurative rhinitis, which was more evident in the mid and caudal turbinates. Epithelial cell cilia loss was observed multifocally at 2 and 4 dpi. At 7 dpi, the same lesions were observed but considered mild. Upon re-inoculation, nasal turbinates showed mild lesions at 23 dpi, that is 2 days post-re-inoculation (2 dpri), and dpi 25 (4 dpri), like those observed on 7 dpi. The amount of SARS-CoV-2 nucleocapsid protein (NP) detected by immunohistochemistry (IHC) in nasal turbinates correlates with the intensity of lesions, showing high, moderate and low amounts of viral antigen at 2 dpi (Supplementary figure 1 (Sp1), a), 4 dpi (Sp1, b) and at 7 dpi (Sp1, c) respectively. Upon re-inoculation, the amount of labelling tended to be mild-to-moderate at 23 dpi (2 dpri) (Sp1, d-e) and very mild at 25 dpi (4 dpri) (Sp1, f-g). Viral antigen was mainly located in nasal epithelial cells, including cells morphologically resembling olfactory neurons and sloughed epithelial cells in the nasal meatus, as well as submucosal gland cells.

Mild non-suppurative tracheitis with very mild patchy loose of the respiratory epithelium cilia was observed at 2 dpi; no evident lesions were observed on the other days. All hamsters tested positive by IHC at 2 dpi, and only two animals had very mild labelling at 4 dpi. No immunohistochemical signal was detected at 7 dpi or after re-inoculation with none of the two SARS-CoV-2 isolates (Supplemental figure 1).

At 2 dpi, all hamsters displayed mild to moderate broncho-interstitial pneumonia (Figure 2a). At 4 dpi, lesions were moderate, including hyperplasia of type II pneumocytes. Multifocal presence of fibrin within alveoli containing inflammatory cells was also observed (Figure 2b). Loss of cilia in affected respiratory airways was evident. The same type of lesions but more severe were observed at 7 dpi (Figure 2c). At 2 dpri and 4 dpri lesions were equivalent to those described at 4 dpi, being from very mild to moderate (Figures 2d and 2e). Immunohistochemical labelling of SARS-CoV-2 NP at 2 dpi varied from low to high and was mainly observed in the bronchial and bronchiolar epithelial cells and type I pneumocytes, following a patchy distribution (mainly in peribronchial and peribronchiolar locations) (Figure 2f). At 4 dpi, SARS-CoV-2 NP protein distribution was similar to that at 2 dpi but with higher amount of labelling in the lung parenchyma and less in the bronchial and bronchiolar epithelia. Noteworthy, type II pneumocytes and mononuclear cells were scarcely labelled (Figure 2g). Lungs tested positive at 7 dpi only in two animals. In these samples, the number of labelled cells was very scarce, and was characterized by the presence of small foci of type I pneumocytes containing viral antigen (Figure 2h). All lung samples were negative at 2 and 4 dpri (respectively 23 and 25 dpi) (Figures 2i and 2j).

**Figure 2.**
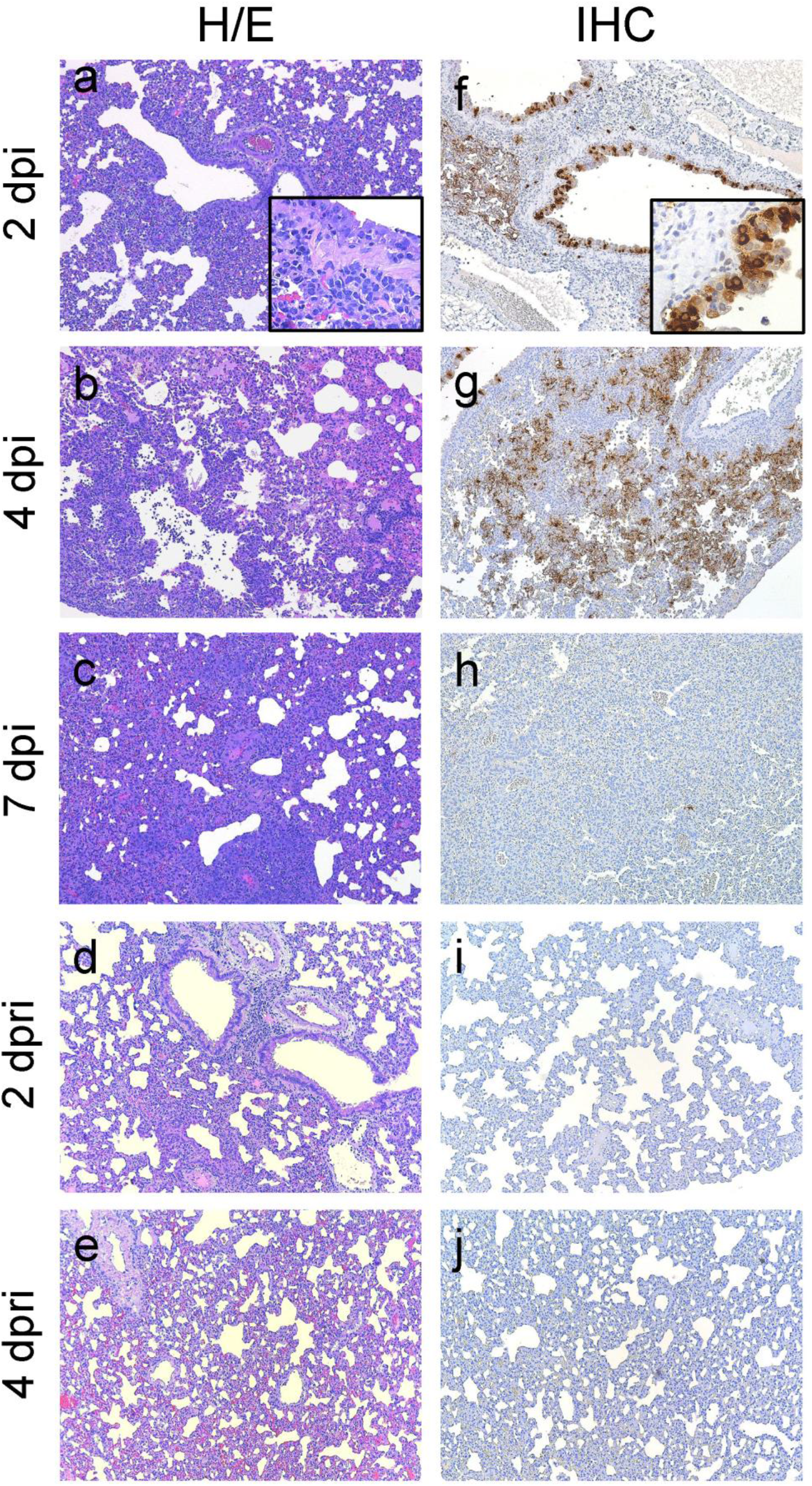
Pathological findings in lungs of hamsters after inoculation and re-inoculation. (a to e) Histopathological findings in lungs of hamsters after SARS-CoV-2 Cat01 challenge on 2 (a), 4 (b) and 7 (c) dpi, and 2 (d) and 4 (e) dpri with Cat01 and WA/1 variants. Broncho-interstitial pneumonia severity increased from 2 to 7 dpi (maximum lesion severity) and was residual at 2 and 4 dpri. Inset in 2a displays submucosa mononuclear inflammation of the bronchus and exocytosis through the epithelium. Hematoxylin and eosin stain, 100x magnification (inset in 2a, 400x magnification) (f to j). Immunohistochemical findings in lungs of same animals. High amount of viral antigen mainly in bronchi epithelium as well moderate amount at 2 dpi (f, inset shows a detail of the bronchus epithelial labelling). The maximum amount of labelling in lung parenchyma, associated to the inflammatory infiltrate, was detected at 4 dpi (g). Scarce number of stained cells were detected at 7 dpi (h, arrowhead) and no labelling was recorded at 2 (i) and 4 (j) dpri. Immunohistochemistry to detect the NP of SARS-CoV-2 and hematoxylin counterstain, 100x magnification (inset in 2a, 400x magnification).

No histopathological findings were found in any of the available mediastinal lymph nodes. By immunohistochemistry, the only available lymph nodes from 2 dpi (n=1) and 4 dpi (n=2), displayed low numbers of dendritic-like cells (stellate appearance) containing viral antigen in the cytoplasm (Supplementary figure 2). No labelling was detected in any of the mediastinal lymph nodes from hamsters at 7 dpi (n=2), 2 dpri (n=3) and 4 dpri (n=4).

No apparent differences in lesion severity or immunohistochemical labelling were observed for any tissue upon re-inoculation with any of the two different SARS-CoV-2 strains. A summary of the severity of lesions and the amount of viral antigen per each individual hamster is displayed in the Supplementary table 1.

### A previous SARS-CoV-2 priming prevents re-infection of the lower respiratory tract

We assessed viral genomic and subgenomic SARS-CoV-2 RNA (gRNA and sgRNA, respectively) levels in nasal turbinates, trachea, lungs and oropharyngeal swabs at 2, 4 and 7 dpi (n=4/day) and 2 and 4 dpri (n=3/day/viral variant) (Figure 3). In addition, we analyzed gRNA and sgRNA levels in oropharyngeal swabs (OS) before the re-challenge to confirm that animals cleared the infection.

**Figure 3.**
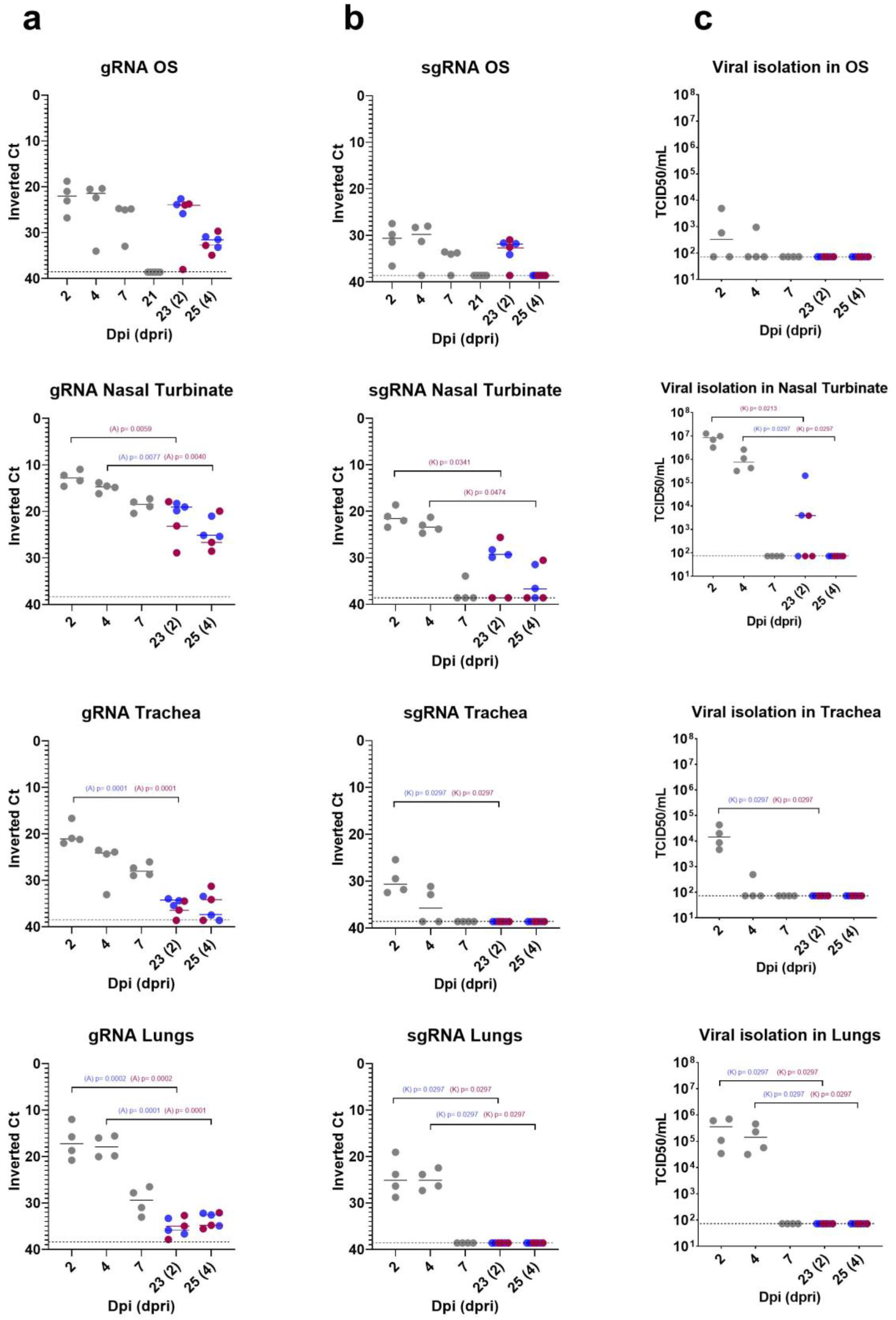
Viral loads in samples obtained from hamsters after inoculation and re-inoculation with SARS-CoV-2. Genomic RNA (a) and subgenomic RNA levels (b) of SARS-CoV-2 was analyzed in oropharyngeal swabs (OS), nasal turbinate, trachea and lungs, as well as the infectious viral loads (c). Horizontal bars reflect median viral loads. In blue data obtained from Cat01-reinfected animals, in red data obtained from WA/1-reinfected animals. Statistically significant p values are reported in the graph, preceded by an (A) for ANOVA test or (K) for Kruskal-Wallis tests.

After the first inoculation, SARS-CoV-2 gRNA loads peaked at 2 dpi in all anatomical compartments and then progressively decreased, accordingly to previous findings ^19^. Viral loads in oropharyngeal swabs were similar at 2 and 4 dpi but decreased by 7 dpi (Figure 3a). Upon re-inoculation, gRNA levels of both viral strains were significantly lower in all sample types in comparison to those obtained during the first infection (Figure 3a). Only limited viral gRNA levels were detected in trachea and lungs of re-infected animals, regardless of the SARS-CoV-2 variant used. we detected higher gRNA levels in Cat01 (G614) re-infected nasal turbinates after 2 dpri once compared to WA/1 (D614) infected samples. This result is in accordance with recent studies ^26^ and suggests an improved viral fitness conferred by the G614 mutation in the upper airway of infected hamsters.

To confirm whether gRNA detection correlated with SARS-CoV-2 replication along the host cells, we monitored the sgRNA levels, following a previously described methodology ^27,28^ (Figure 3b). The levels of SARS-CoV-2 sgRNA were lower than gRNA levels in all hamster samples but also peaked at 2dpi and followed a similar declining trend over time, as previously reported ^14,27^. Detection of sgRNA at 7 dpi was only possible in one nasal turbinate and three out of four oropharyngeal swab samples. Once the animals were re-inoculated, sgRNA of both viral variants was detected in nasal turbinates at 2 and 4 dpri, but only at 2 dpri in OSs. We did not detect sgRNA in trachea and lungs at any time point post-re-inoculation, suggesting that both viral strains were not able to successfully replicate in these organs upon reinfection (Figure 3b).

Then, we compared the SARS-CoV-2 gRNA and sgRNA loads obtained from the two infection events. We found statistically significant reductions of both genomic and subgenomic viral RNA levels in samples of reinfected animals when compared to those obtained after the first challenge; statistics are summarized in Supplementary table 2. In addition to the molecular detection of gRNA and sgRNA, the infectious viral titers were measured in Vero E6 cells and expressed as TCID_50_/mL. We detected infectious virus particles in all tissues and OSs at 2 and 4 dpi; however, no infectious virus samples were found from 7 dpi onwards (Figure 3c). Infectious viral loads were particularly high in nasal turbinates at 2 dpi and 4 dpi. The lack of virus isolation at 7 dpi samples (consistent with the low levels of sgRNA and viral antigen detection) and the remission of clinical signs of animals (weight) starting from 5 dpi, indicate that these animals possibly cleared the infection and stopped shedding infectious virus around 7 dpi.

After re-challenge, at 2 dpri, we recorded the presence of infectious virus in two out of four nasal turbinate samples of hamsters re-exposed to the Cat01 isolate. Similarly, only one animal inoculated with the WA/1 variant displayed infectious virus in the nasal turbinates. Re-infected positive animals were much less susceptible to infection than those experimenting a first challenge (Figure 3c). Indeed, values’ variations obtained by quantifying genomic RNA, subgenomic RNA or infectious viral particles were statistically significant (Supplementary Table 2). Conversely, we were not able to identify infectious SARS-CoV-2 particles from trachea, lungs or OSs of re-inoculated hamsters at any time point, regardless of the viral variant used.

### Hamsters developed a neutralizing humoral immune response against SARS-CoV-2 from 7 days post infection

To monitor humoral immune responses elicited by golden Syrian hamsters upon infection with SARS-CoV-2, we used the serum of individual hamsters collected until 7 dpi and those generated during the re-infection period (2 and 4 dpri). In addition, serum was prepared from animals before the reinfection procedure (21 dpi) and pooled in groups of four (according to animal caging at the BSL3 facility) to detect the levels of antigen-specific humoral response against SARS-CoV-2. The levels of immunoglobulins targeting specific viral antigens encompassing the S protein (S1+S2), the receptor binding domain (RBD) and the NP were detected using an *in house* ELISA (Figure 4a, 4b and 4c). Seroconversion against all tested proteins was evident by 7 dpi, although S specific antibody levels were notably lower than those against the RBD and NP, probably due to a lower sensitivity of the S-ELISA compared with the RBD one. Data obtained from sera collected at 21 dpi suggested that total antibody levels targeting the S glycoprotein and those recognizing specifically the RBD subdomain incremented between 7 and 21 dpi. Conversely, the level of NP-specific antibodies notably decreased at 21 dpi. After re-inoculation, the levels of antibodies against S, RBD and NP antigens further increased until 25 dpi (4 dpri) independently of the strain.

**Figure 4.**
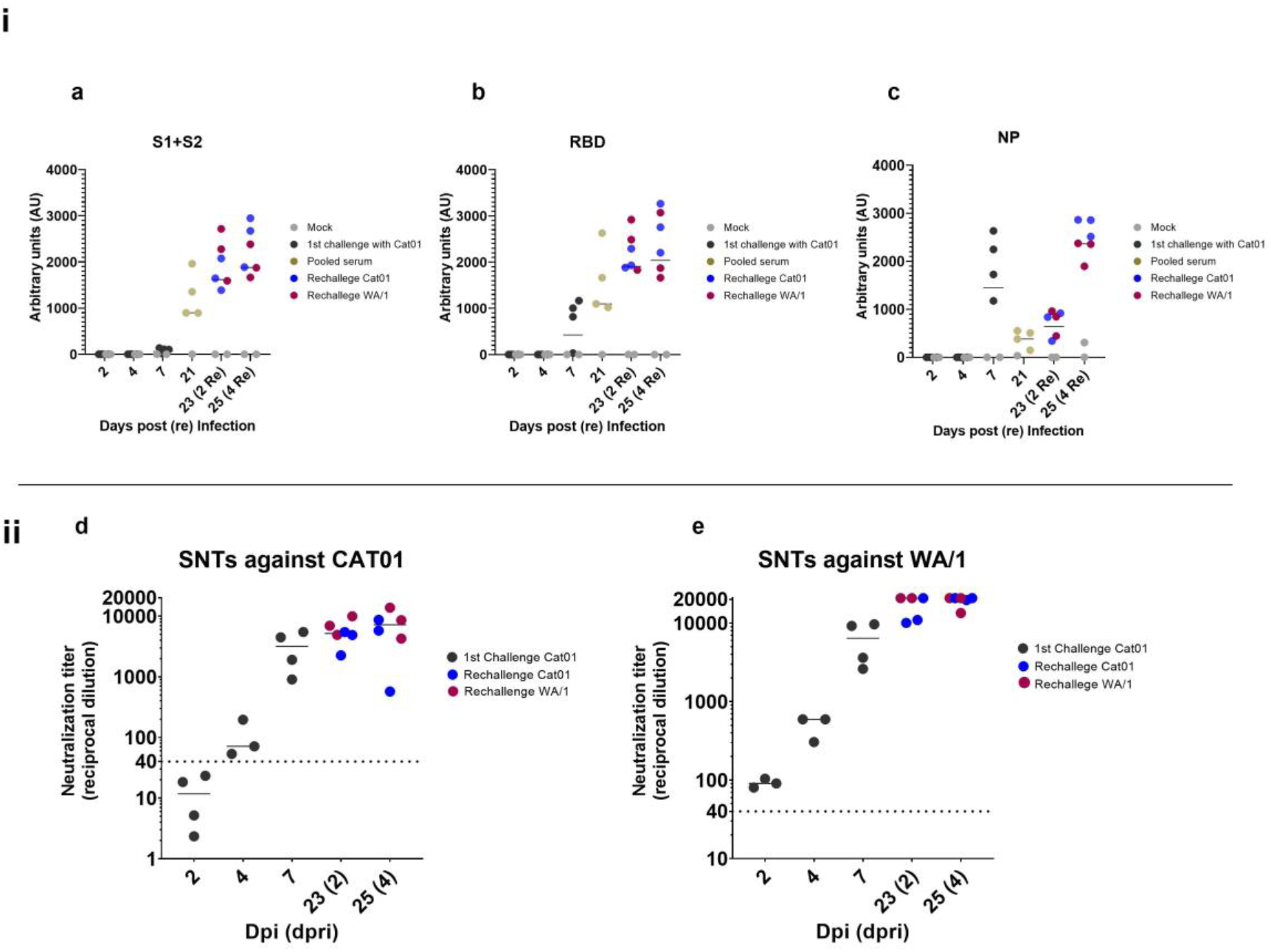
Humoral responses in SARS-CoV-2 reinfected hamster. i) Antibody subclasses against a) Spike protein subunits 1 and 2, b) receptor binding domain (RBD) and c) nucleocapsid protein. In black, serum samples from animals challenged with Cat01 (1st inoculum), in blue serum from animals re-inoculated with the same viral variant (Cat01), in red serum from animal re-inoculated with the different viral variant (WA/1). At 21 dpi before the re-inoculation sera were collected from Cat01-exposed animals, pooled following housing criteria and analyzed (in gold). In grey, serum from control animals treated with mock solution of PBS. ii) Serum from all animals were used for live virus neutralization assay against d) Cat01 and e) WA/1 variants. Code color are the same that those used in panel i). Pooled sera were not tested in the SNT assay.

We then evaluated the neutralization activity of sera obtained from all animals excepted those collected at 21 dpi. Neutralization activity was detected from 7 dpi onwards, and sharply increased at 2 and 4 dpri (Figures 4d and 4e), similar to kinetic profiles of antigen-specific antibodies detected by ELISA. Importantly, the humoral response generated against the Cat01 strain after the primary infection was able to neutralize G614 and D614 viral isolates (Cat01 and WA/1, respectively). Moreover, animals were protected from reinfection at the lower respiratory tract against both homologous and heterologous variants. Thus, the high titer of neutralizing antibodies detected already at 7 dpi (and onwards) might explain the partial protection (at the lower respiratory tract but not the upper one) of these animals against the re-challenge, which further caused a booster neutralizing effect.

## Discussion

The golden Syrian hamster represents an animal model able to recapitulate some infection/re-infection events found in humans. Specifically, this model is characterized by mild-to-moderate clinical, pathological and virological outcomes upon infection with SARS-CoV-2, and minimal virological outcome at the upper respiratory tract upon reinfection with a D164-homologous or a G614-heterologous SARS-CoV-2 variants.

Results obtained during the primary infection are comparable to those reported in previous studies ^13,18,19^, confirming the natural susceptibility of this animal model to SARS-CoV-2. The golden Syrian hamster reproduces the mild-to-moderate COVID-19 induced pathology described in humans, especially in the lower respiratory tract ^20^. Furthermore, we also demonstrated for the first time that reinfection can occur at the upper respiratory tract of this species using either homologous or heterologous SARS-CoV-2 variants. However, our results of virus titration, gRNA and sgRNA detection indicate that negligible or nil viral infection and replication occurred in the lower respiratory tract of re-infected hamsters, regardless of the viral variant used. Conversely, both viral isolates were able to infect nasal turbinates of animals having resolved a primary infection. These results are aligned with those described by van Doremalen *et al.* ^29^, which showed that immune responses generated by an adenovirus-vectorized ChAdOx1 vaccine, encoding the spike protein of SARS-CoV-2, prevented viral replication in the lower respiratory track of challenged rhesus macaques but not at the nasal turbinates. Importantly, no viral gRNA reduction was recorded in the upper respiratory tract of non-human primates and vaccinated animals shed high quantities of infectious virus. Similarly, we found infectious SARS-CoV-2 particles in 1 or 2 out of 4 nasal turbinate samples of hamsters re-exposed to the SARS-CoV-2 WA1 or CAT01 isolates, respectively. These results suggest that human re-infections might also occur in the upper respiratory tract since a previously generated immunity is not sterilizing at the nasal level.

Our study showed that infected animals developed specific humoral immune responses against different proteins of the SARS-CoV-2 from 7 dpi onwards. As mentioned before, S and RBD specific antibodies increased after 7dpi. Conversely, levels of NP-specific antibodies decreased until day 21 pi. In a recent study, *Chen et al.* ^30^ analyzed the IgG kinetic profile of infected human patients, highlighting that SARS-CoV-2 infection induced a distinct temporal profile of humoral responses against different viral proteins. These observations demonstrated that levels of NP-specific IgGs tend to sharply decrease after 14-20 days post-symptoms in humans, similar to our observation in the hamster model. Moreover, the virus neutralization assay indicated that infected hamsters developed high levels of SARS-CoV-2 neutralizing antibodies. Previous studies in COVID-19 patients have correlated levels of serological and mucosal neutralizing antibodies ^31,32^, which have also been associated to infection clearance. The same phenomenon could have occurred in the golden Syrian hamster model after the primary SARS-CoV-2 infection. These animals might have mounted a strong mucosal and serological humoral response able to neutralize the virus at the lower respiratory tract upon a secondary infection, as evidenced by the high seroneutralizing antibody responses from 7 dpi onwards. This protective effect was observed in trachea and lungs independently of the viral variant used for re-inoculation purposes. Remarkably, the immunity developed by challenged hamsters was not sterilizing, and localized infection at the upper respiratory tract was still possible upon reinfection. However, animals rapidly cleared the virus by day 4 dpri, similarly to what was previously described in African Green monkeys ^33^ and Rhesus macaques ^15^ models.

In line with results obtained using the golden Syrian hamster model, protection from a secondary infection was also described in rhesus macaques after two consecutive SARS-CoV-2/WA1 inoculations ^14^. Although low levels of viral replication were reported in the nasal cavity of these non-human primates, disease prevention was also associated with the presence of high neutralizing antibody levels in sera ^14^, a fact that would also apply in the studied hamsters.

On the other hand, previous studies in human patients have characterized T cell responses against SARS-CoV and SARS-CoV-2 ^34,35^, demonstrating the important role of the cellular components of the immunity to resolve these Sarbecovirus infections. In addition, specific T cell responses generated by the pre-exposition to seasonal human coronaviruses (HCoV)-229E and OC43 or the zoonotic SARS-CoV have been demonstrated to cross-react and also target SARS-CoV-2 epitopes ^36,37^. A recent study from *Brocato et al.* ^13^ proved that disruption of lymphocytes in immunosuppressed Golden Syrian hamsters severely affected the clinical outcome of the SARS-CoV-2 infection, resulting in an increased weight loss and higher viral loads found in the lungs, and suggesting a critical role of T-cells in the early phase of infection. The role of cellular and mucosal immunity was not addressed in the current work and, therefore, further studies would be required to understand immune mechanisms involved in the protection against SARS-CoV2 reinfection.

It was already known that seasonal HCoVs like HKU1, OC43, or 229E can re-infect the same host ^38,39^, suggesting a similar potential scenario for SARS-CoV-2. In fact, several confirmed cases of SARS-CoV-2 reinfection have been described worldwide since mid-August 2020 ^7–11^. To date, all reinfection cases were described by sequencing different SARS-CoV-2 variants between the first and second episodes of infection. In the golden Syrian hamster model both homologous and heterologous variant reinfections were possible. Re-exposed animals did not show evident clinical signs during the short phase of reinfection (2 to 4 days). Conversely, reinfections in humans have been linked to different clinical outcomes. Symptomatic reinfection ranged from more severe ^8,10^ to milder forms ^7 9^. Furthermore, asymptomatic reinfection has also been described ^11^. This variability might be the result of multiple factors as the overall health condition of the patient, the initial viral load or reinfection with a viral strain with higher replication fitness and, importantly, the level of neutralizing antibodies and specific T-cells at the time of subsequent infection event. More studies are needed to establish if different clinical outcomes are possible in case of reinfection using the hamster model. This might be obtained by modifying several aspects of the experimental design as for example increasing the viral load between challenges or using immunosuppressed animals as suggested by Brocato *et al* ^13^.

The overall frequency of reinfections in the human population is difficult to establish, since a high number of them could pass unnoticed. These could lead to underestimate the rate of reinfection cases, especially if the period between infections was particularly short. In fact, a recent report highlighted that reinfection is possible within one month from the onset of clinical symptoms ^10^. Currently, the incidence rate of reinfection is estimated to be approximately 0.36 per 10,000 person/week ^12^, which is rather significant numerically, taking into account the high number of new infection cases being reported daily ^1^. Therefore, massive testing and systematic monitoring of the population that has been infected once would probably lead to a much higher detection of reinfections than those reported so far. In our study, the percentage of reinfected animals showing infectious virus in the upper respiratory tract was notable: 1 out of 3 with animals receiving the heterologous variant and 2 out of 3 that were re-inoculated with the homologous variant. These data emphasize the interest of the golden Syrian hamster as a model to study reinfection mechanisms.

In summary, the golden Syrian hamster can be considered a convenient animal model to study reinfection pathogenesis and immune responses. Specifically, this animal species can suffer from a clinical frame resembling mild-to-moderate disease in humans, and reinfection with homologous (G614) or heterologous (D614) variants yielded to a subclinical infection of the upper respiratory tract. No sterilizing immunity was elicited after the first infection event; however, the lower respiratory tract was fully protected upon re-inoculation with both viral variants

## Methods

### Ethics statement

Animal experiments were approved by the Institutional Animal Welfare Committee of the Institut de Recerca i Tecnologia Agroalimentàries (CEEA-IRTA, registration number CEEA 188/2020) and by the Ethical Commission of Animal Experimentation of the Autonomous Government of Catalonia (registration number FUE-2020-01589810) and conducted by certified staff. Experiments with SARS-CoV-2 were performed at the Biosafety Level-3 (BSL-3) facilities of the Biocontainment Unit of IRTA-CReSA (Barcelona, Spain).

### Virus isolates

Two different SARS-CoV-2 isolates were used: hCoV-19/Spain/CT-2020030095/2020 (GISAID ID EPI_ISL_510689), designated as Cat01, and hCoV-19/USA/WA1/2020 (GISAID ID EPI_ISL_404895), designated as WA/1. The WA/1 isolate was kindly provided by Dr. Slobodan Paessler (University of Texas, USA). Cat01 was isolated from human patient (Oropharyngeal swab) from Spain in March 2020.

Compared to Wuhan/Hu-1/2019 strain, Cat01 isolate has the following point mutations: D614G (Spike), R682L (Spike), C16X (NSP13) and other 12 in NSP3 (M1376X, P1377X, T1378X, T1379X, I1380X, A1381X, K1382X, N1383X, T1384X, V1385X, K1386X, S1387X).

SARS-CoV-2 WA1 was isolated from a human patient (Oropharyngeal swab) from Washington State (US) in January 2020 and differs from Wuhan/Hu-1/2019 strain for the presence of a single point mutation: L84S (NS8).

Production of virus stocks (Cat01 passage number 3; WA/1 passage number 2), isolation, titration and live virus neutralization assay were performed in Vero E6 cell (ATCC® CRL-1586™). Virus titers were determined using a standard TCID_50_ assay and expressed as TCID_50_ /mL.

### Study design

A total of thirty-four 5-6-week-old male and female golden Syrian hamsters (Charles River) were used in this study (Figure 5). Under isoflurane anesthesia, 24 hamsters were inoculated by intranasal instillation with 10^5.8^ TCID_50_ of the SARS-CoV-2 Cat01 isolate per animal (100 μL/individual, 50 μL for each nostril). Ten hamsters were mock intranasally inoculated with PBS (100 μL/individual, 50 μL for each nostril) and used as negative controls. Body weight was monitored daily during the first week post-inoculation and 4 days post-re-inoculation. For virological and pathological examinations, 4 inoculated and 2 control hamsters (half male and half female) were sacrificed on days 2, 4 and 7 dpi. The weight of remaining hamsters (n= 12 inoculated and n= 4 control) was recorded at 10, 14 and 21 dpi.

**Figure 5.**
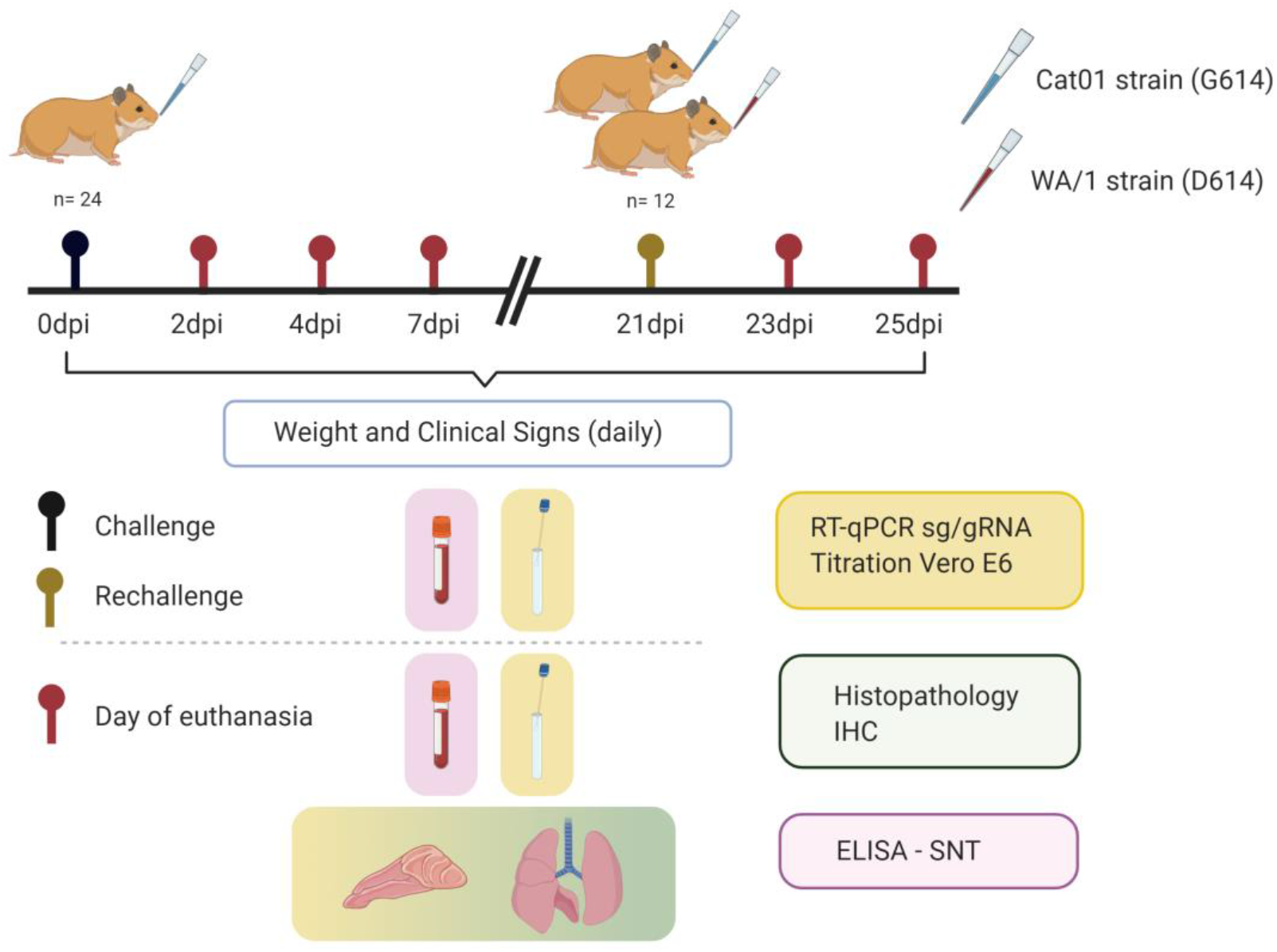
Experimental Design. Golden Syrian hamsters (n=24, 12 male and 12 female) were intranasally (IN) inoculated with 10^5.8^ TCID_50_ of SARS-CoV-2 Cat01 isolate. Before challenge blood samples and oropharyngeal swabs (OS) were collected from all animals. At 2-, 4- and 7-days post-inoculation (dpi), 4 infected animals (2 male and 2 female) were euthanized. Before necropsy, blood samples and OS were collected from each animal. Nasal turbinate, lungs and trachea were collected for pathological and virological analyses. In purple, samples used for ELISA and seroneutralization test; in green, samples used for histopathology and immunohistochemistry; in yellow, samples used for RT-qPCR and viral titration in Vero E6 cell. At 21 dpi, the remaining animals (n = 12) were equally divided in two experimental groups. One group was intranasally inoculated with 10^5.2^ TCID_50_ of Cat01 isolate while the other was IN inoculated with WA/1 strain at the same concentration. At day 23-dpi (2 days-post re-inoculation) and 25-dpi (4 days-post re-inoculation), 3 animals/experimental group were euthanized. Sampling was equivalent to that indicated previously. Created with BioRender.com

At 21 dpi, 6 animals from the inoculated group were intranasally re-inoculated with 10^5.2^ TCID_50_ of SARS-CoV-2 Cat01 isolate (100 μL/individual, 50 μL for each nostril), while the other 6 animals received the same dose of the WA1 isolate. The four control hamsters were re-inoculated with the same amount of PBS serving as negative controls of the re-inoculation. Inoculation procedures were the same as those used for the first part of the experiment. For virological and pathological examinations, 3 inoculated animals for each viral isolate and 2 control animals were sacrificed at 23 dpi (2 dpri) and 25 dpi (4 dpri). At necropsy, samples from nasal turbinate, trachea and lung were taken and fixed by immersion in 10% buffered formalin. For molecular detection and viral titration purposes, a portion of the same tissues were placed in individual Eppendorf tubes containing 500 μl of DMEM (GIBCO) supplemented with 1% penicillin-streptomycin (PS) (GIBCO) with a single zinc-plated, steel, 4.5-mm bead. Samples were homogenized at 30 Hz for 2 min using a TissueLyser II (QIAGEN GmbH, Hilden, Germany) and centrifuged for 30 sec at 11,000 rpm. All samples were stored at −70ºC. Blood samples were collected from each animal at the necropsy day by cardiac puncture under deep anesthesia, centrifugated 10 minutes at 2100 rpm at RT and stored at −70ºC for further analysis. Similarly, oropharyngeal and rectal swabs were collected before the sacrifice under deep anesthesia, resuspended in 500 μl of DMEM supplemented with 1% PS and stored at −70ºC for further analysis. Clinical evolution as well as pathological, immunological and virological outcomes were evaluated at different time points as indicated in Figure 5.

### Pathology and immunohistochemistry

Upper (nasal turbinate) and lower (trachea and lung) respiratory tract formalin-fixed samples were routinely processed for histopathology and hematoxylin&eosin stained slides examined under optical microscope. The mediastinal lymph node was also studied in 13 out of the 34 analyzed animals, since not in all cases was identified while taking samples during necropsy or during the trimming of tissues. A semi-quantitative approach based on the amount of inflammation (none, mild, moderate, or severe) was used to score the damage caused by SARS-CoV-2 infection in hamsters.

A previously described immunohistochemistry technique to detect SARS-CoV-2 NP antigen ^40^ using the rabbit monoclonal antibody (40143-R019, Sino Biological, Beijing, China) at dilution 1:1000, was applied on nasal turbinates, trachea, lung and mediastinal lymph nodes. The amount of viral antigen in tissues was semi-quantitatively scored in the different studied tissues (low, moderate and high amount, or lack of antigen detection).

### RNA-extraction and quantitative RT-PCR

Viral RNA was extracted from target organs and swabs samples using the IndiMag pathogen kit (Indical Bioscience) on a Biosprint 96 workstation (QIAGEN) according to the manufacturer’s instructions. RT-PCR used to detect viral gRNA is based on the one published by Corman *et al.* (2020), with minor modification to adapt it to the AgPath-ID One-Step RT-PCR Kit (Life Technologies). RT-PCR targets a portion of envelope protein gene (position 26,141-26,253 of GenBank NC_004718). The primers and probes used, and their final concentration are the follow: forward: 5’–ACAGGTACGTTAATAGTTAATAGCGT – 3’ [400 nM], reverse: 5’-ATATTGCAGCAGTACGCACACA –3’ [400 nM] probe: 5’-FAM-ACACTAGCCATCCTTA CTGCGCTTCG-TAMRA – 3’ [200 nM]. Thermal cycling was performed at 55°C for 10 min for reverse transcription, followed by 95°C for 3 min and then 45 cycles of 94°C for 15 s, 58°C for 30 s. sgRNA detection by RT-PCR is based on protocol published by Wölfel *et al.* (2020) with minor modification to adapt it to the AgPath-ID One-Step RT-PCR Kit. The primers and probes are the same used for gRNA detection except for primer forward: 5’-CGATCTCTTGTAGATCTGTTCTC –3’ [400 nM]. [400 nM]. Thermal cycling was performed at 55°C for 10 min for reverse transcription, followed by 95°C for 3 min and then 45 cycles of 95°C for 15 s, 56°C for 30 s.

### Evaluation of the humoral response against SARS-CoV-2 by ELISA

The level of anti-SARS-CoV-2 antibodies in hamster serum samples was determined using a sandwich-ELISA. Nunc MaxiSorp ELISA plates were coated overnight at 4ºC with 50 μL of capture antibody (anti-6xHis antibody, clone HIS.H8; ThermoFisher Scientific) at 2 μg/mL in PBS. Then, plates were blocked for two hours at room temperature using PBS/1% of bovine serum albumin (BSA) (Miltenyi biotech) and 50 μL (1μg/mL in blocking buffer) of the SARS-CoV-2 Spike (S1+S2), receptor-binding domain (RBD) or nucleocapsid protein (NP) (Sino Biologicals, Beijing, China) were added and incubated overnight at 4ºC. Each sample was diluted in blocking buffer and assayed in duplicated. Diluted samples were incubated overnight at 4ºC. Antigen free wells were also assayed in parallel in the same plate to evaluate sample background. Serial dilutions of a positive serum sample were used as standard. As secondary antibody, an HRP-conjugated Goat anti-hamster IgG (H+L) (Jackson Immunoresearch) at 1/20,000 dilution in blocking buffer was used. Secondary antibody was incubated for one hour at room temperature. Plates were revealed using o-Phenylenediamine dihydrochloride (OPD) (Sigma Aldrich) and stopped with 4N of H2SO4 (Sigma Aldrich). The signal was analyzed as the optical density (OD) at 492 nm with noise correction at 620 nm. The specific signal for each antigen was calculated after subtracting the background signal obtained for each sample in antigen-free wells. Results are shown as arbitrary units (AU).

### Live virus neutralization assay

Prior neutralization assay, sera samples were heat inactivated at 56 ºC for 30 minutes. Inactivated sera samples were serially 2-fold diluted (range 1/40 to 1/20,480) in DMEM supplemented with 100 U/mL penicillin, 100 μg/mL streptomycin, and 2 mM glutamine (all ThermoFisher Scientific), mixed with SARS-CoV-2 Cat01 or WA/1 isolates and further incubated at 37 C for 1 hour). Each dilution (in duplicates) containing 100 TCID of virus solution. The mixtures were then transferred to Vero E6 cell monolayers (ATCC CRL-1586) and cultured for 3 days at 37 C and 5% CO_2_. Cytopathic effects of the virus were measured after 3 days using the CellTiter-Glo luminescent cell viability assay (Promega), according to the manufacturer’s protocol. Luminescence was measured in a Fluoroskan Ascent FL luminometer (ThermoFisher Scientific).

### Statistical analysis

Statistical analyses were completed using GraphPad Prism 8. Mean weight data were analyzed using a multiple unpaired *t*-test for comparison of 2 groups (infection) and Mixed-effect model (REML) with Dunnett’s multiple comparisons test for the comparison of 3 groups (reinfection). Normality of each data set for gRNA, sgRNA and infection viral loads was calculated using Shapiro-Wilk test. Comparison of viral loads between primary challenge and rechallenge was performed with ordinary one-way ANOVA and Dunnett’s multiple comparison (paired test) or Kruskal-Wallis and Dunn’s multiple comparison test (unpaired test). In all analyses, a P value <0.05 was considered statistically significant.

## Author contributions

MB contributed to perform experiments, analyze data and write the manuscript; JR contributed to perform experiments, analyze data and write the manuscript; MLRC contributed to serological studies; CAV contributed to serological studies; GC contributed to perform animal experiment; MP performed histopathology and IHC of samples; NT contributed to analyze data and prepare figures; MNJ contributed to molecular biology studies; VG contributed to project conceptualization; AV contributed to project conceptualization; NR contributed to laboratory work; NIU contributed to project conceptualization; JB contributed to project conceptualization; BC contributed to project conceptualization and funding; AB contributed to project conceptualization; JC contributed to serological studies and project conceptualization; JVA contributed to design and perform the experiment, analyze the data, funding and prepare the manuscript; JS contributed to design and perform the experiment, analyze the data, funding and prepare the manuscript. All authors have revised and approved the manuscript.

