## Supplementary Figures and Tables for "Protection against reinfection with D614- or G614-SARS-CoV-2 isolates in hamsters"

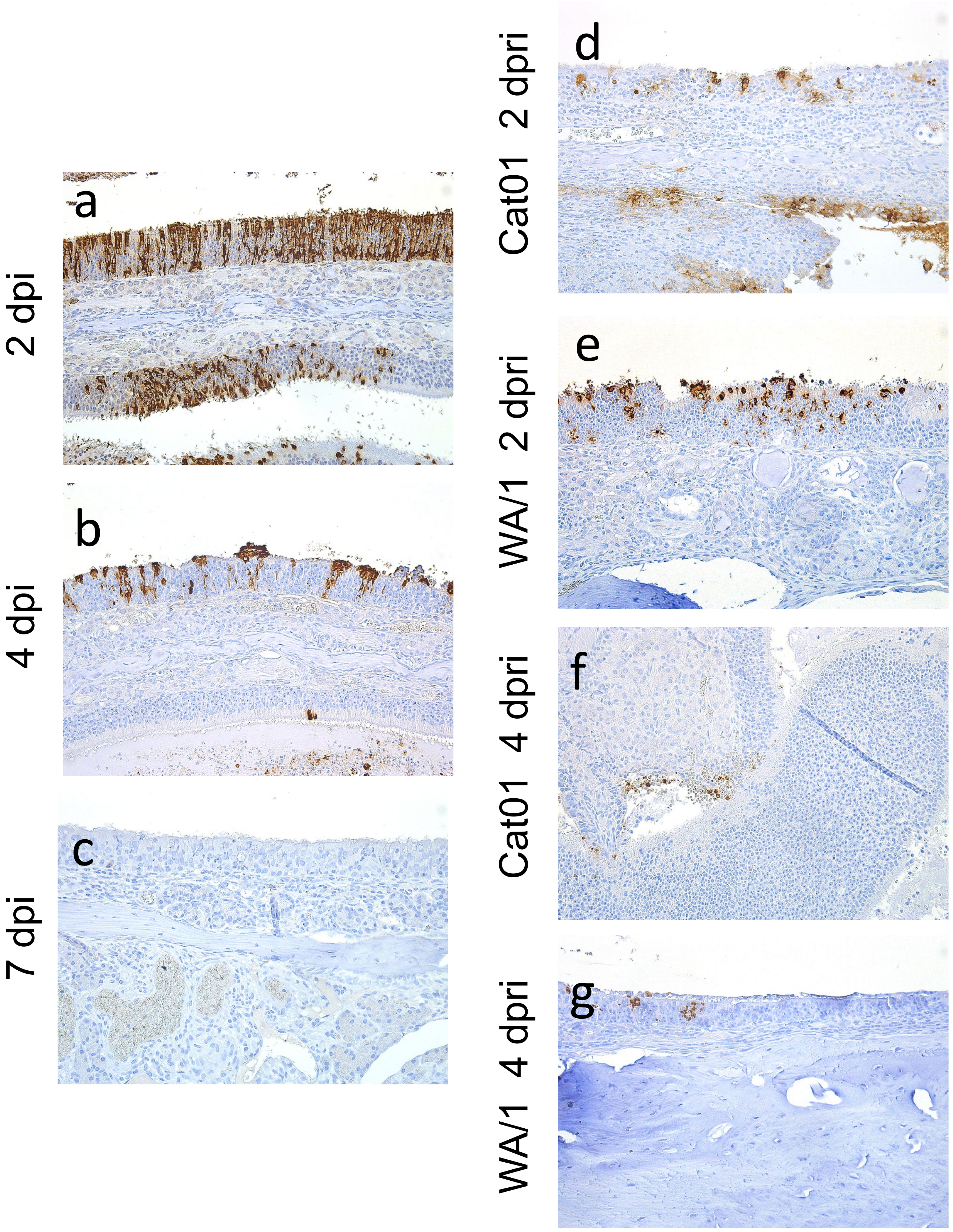
a
2 dpi
b
4 dpi
c
7 dpi
d
Cat01 2 dpri
e
WA/1 2 dpri
f
Cat01 4 dpri
g
WA/1 4 dpri

**Supplementary figure 1. Immunohistochemical detection of SARS-CoV-2 in nasal turbinates.** Detection of viral antigen in nasal turbinates of hamsters inoculated with SARS-CoV-2 Cat01 isolate and euthanized on days post-inoculation (dpi) 2 (a), 4 (b) and 7 (c) and days post-re-inoculation (dpri) 2 (d,e) and 4 (f,g) with Cat01 and WA/1 isolates. The highest amount of viral antigen was in nasal epithelial cells at 2 dpi (a), decreasing significantly on dpi 4 (b). No viral antigen was found on dpi 7 (c). Moderate amount of labelling was found on dpri 2 following re-inoculation with both isolates (d,e) and was low by dpri 4 (f,g). Immunohistochemistry to detect the NP of SARS-CoV-2 and hematoxylin counterstain, 100x magnification.

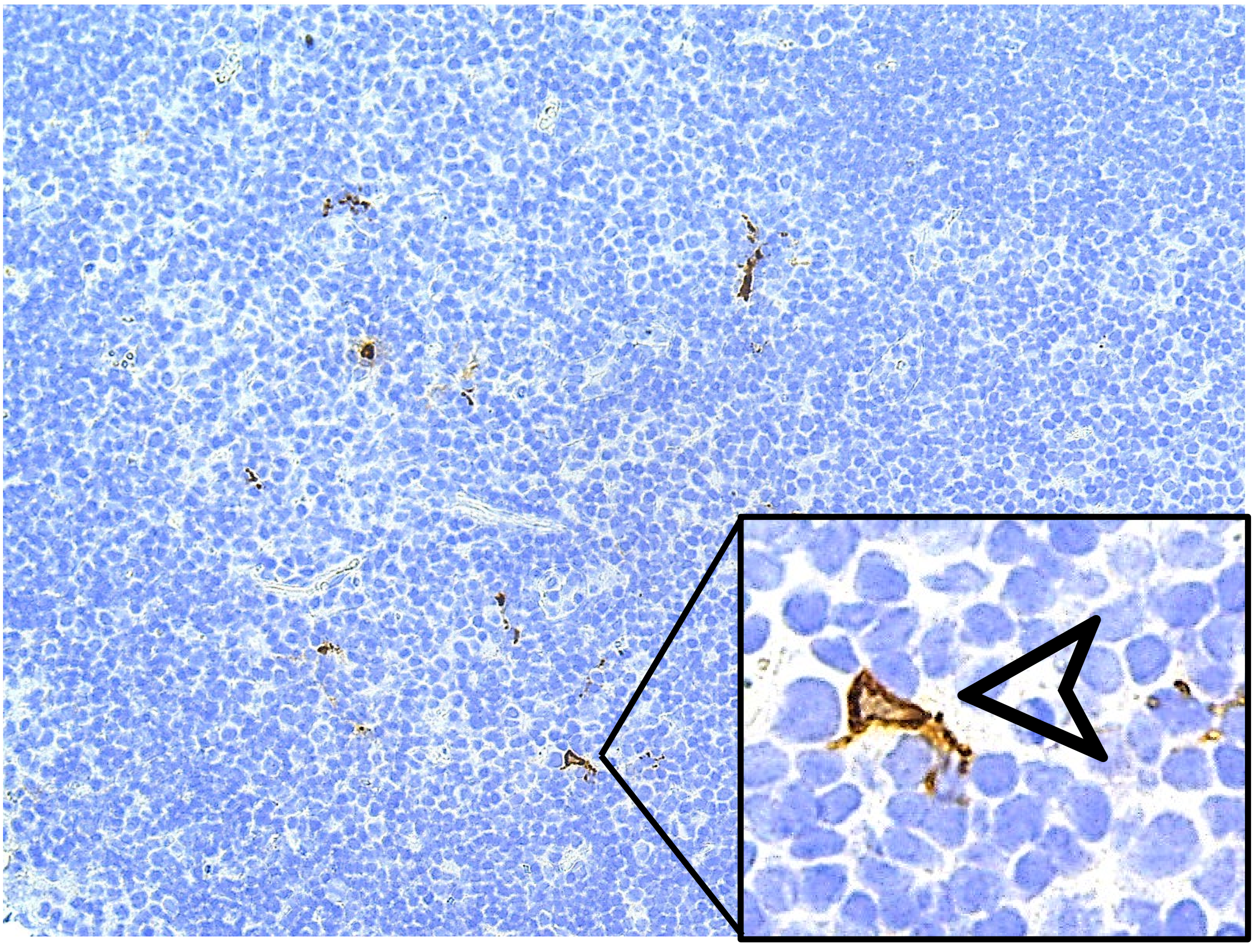

**Supplementary figure 2. Immunohistochemical detection of SARS-CoV-2 in mediastinal lymph node.** Detection of viral antigen in the cytoplasm of scattered dendritic-like cells in a histologically normal mediastinal lymph node of a hamster inoculated with SARS-CoV-2 Cat01 isolate at dpi 2. Inset: magnification of one dendritic-like cells showing labelling in the cytoplasm and dendritic prolongations. Immunohistochemistry to detect the NP of SARS-CoV-2 and hematoxylin counterstain, 100x magnification (inset, 400x magnification).

**Supplementary table 1**. Summary of the histopathological findings (HIST) in nasal turbinate, trachea, lung and mediastinal lymph node (LN), as well as the immunohistochemical (IHC) results of the same tissues, per individual studied hamster.

| Hamster ID | dpi/dpri | HIST Nasal turbinate | IHC Nasal turbinate | HIST Trachea | IHC Trachea | HIST Lung | IHC Lung | HIST Mediast. LN | IHC Mediast. LN |
|---|---|---|---|---|---|---|---|---|---|
| 1 | 2 | ++ | +++ | + | + | + | ++ | NA | NA |
| 2 | 2 | +++ | +++ | + | + | ++ | +++ | NA | NA |
| 5 | 2 | + | +++ | - | + | ++ | +++ | - | + |
| 6 | 2 | +++ | +++ | + | + | + | + | NA | NA |
| 25* | 2 | - | - | - | - | - | - | NA | NA |
| 27* | 2 | - | - | - | - | - | - | NA | NA |
| 3 | 4 | ++ | ++ | - | - | ++ | ++ | NA | NA |
| 4 | 4 | +++ | + | - | - | ++ | ++ | NA | NA |
| 7 | 4 | ++ | ++ | - | + | ++ | +++ | - | + |
| 8 | 4 | ++ | ++ | - | + | ++ | +++ | - | + |
| 26* | 4 | - | - | - | - | - | - | NA | NA |
| 28* | 4 | - | - | - | - | - | - | NA | NA |
| 9 | 7 | + | + | - | - | ++ | - | - | - |
| 10 | 7 | + | - | - | - | +++ | - | NA | NA |
| 13 | 7 | + | + | - | - | +++ | + | - | - |
| 14 | 7 | + | + | - | - | ++ | + | NA | NA |
| 29* | 7 | - | - | - | - | - | - | NA | NA |
| 33* | 7 | - | - | - | - | - | - | - | - |
| 11 | 23/2 | + | ++ | - | - | ++ | - | NA | NA |
| 12 | 23/2 | + | - | - | - | ++ | - | - | - |
| 15 | 23/2 | + | + | - | - | + | - | NA | NA |
| 16 | 23/2 | ++ | ++ | - | - | + | - | - | - |
| 20 | 23/2 | + | ++ | - | - | + | - | NA | NA |
| 24 | 23/2 | ++ | ++ | - | - | + | - | - | - |
| 30* | 23/2 | + | - | - | - | - | - | NA | NA |

| | | | | | | | | | |
|---|---|---|---|---|---|---|---|---|---|
| 34* | 23/2 | - | - | - | - | - | - | NA | NA |
| 17 | 25/4 | + | - | - | - | + | - | NA | NA |
| 18 | 25/4 | + | + | - | - | + | - | - | - |
| 19 | 25/4 | + | + | - | - | + | - | - | - |
| 21 | 25/4 | - | - | - | - | + | - | NA | NA |
| 22 | 25/4 | + | + | - | - | + | - | - | - |
| 23 | 25/4 | + | + | - | - | + | - | - | - |
| 31* | 25/4 | - | - | - | - | - | - | NA | NA |
| 35* | 25/4 | - | - | - | - | - | - | NA | NA |

HIST: histopathological findings (-, no lesions; +, mild lesions; ++, moderate lesions; +++, severe lesions)

IHC: immunohistochemical labelling (-, none; +, low amount; ++, moderate amount; +++, high amount)

NA: non-available

* Non-SARS-CoV-2 inoculated hamsters (negative control animals)

| | Cat01-Cat01 | | Cat01-WA/1 | |
|---|---|---|---|---|
| | 2-dpi / 2-dpri | 4-dpi / 4-dpri | 2-dpi / 2-dpri | 4-dpi / 4-dpri |
| | genomic RNA | | | |
| Oralpharyngeal Swab | ns | ns | ns | ns |
| Nasal Turbinate | ns | (A) *p*=0.0077 | (A) *p*=0.0059 | (A) *p*=0.0040 |
| Trachea | (A) *p*=0.0001 | ns | (A) *p*=0.0001 | ns |
| Lungs | (A) *p*=0.0002 | (A) *p*=0.0001 | (A) *p*=0.0002 | (A) *p*=0.0001 |
| | subgenomic RNA | | | |
| Oralpharyngeal Swab | ns | ns | ns | ns |
| Nasal Turbinate | ns | ns | (K) *p*=0.0341 | (K) *p*=0.0474 |
| Trachea | (K) *p*=0.0297 | ns | (K) *p*=0.0297 | ns |
| Lungs | (K) *p*=0.0297 | (K) *p*=0.0297 | (K) *p*=0.0297 | (K) *p*=0.0297 |
| | Viral titration | | | |
| Oralpharyngeal Swab | ns | ns | ns | ns |
| Nasal Turbinate | ns | ns | (K) *p*=0.0213 | (K) *p*=0.0474 |
| Trachea | (K) *p*=0.0297 | ns | (K) *p*=0.0297 | ns |
| Lungs | (K) *p*=0.0297 | (K) *p*=0.0297 | (K) *p*=0.0297 | (K) *p*=0.0297 |

**Supplementary table 2. Genomic RNA, subgenomic RNA and infectious viral loads comparison between infection and reinfection.** Viral loads obtained at 2 and 4 dpi were compared to those obtained at 2 and 4 dpri, in case of homologous (Cat01-Cat01) or heterologous (Cat01-WA/1) variant re-inoculation. P values indicate a statistically significant reduction in viral loads of re-inoculated animals. ns= not statistically significant. Comparisons between primary challenge and rechallenge viral loads were performed with ordinary one-way ANOVA and Dunnett's multiple comparison (A) or Kruskal-Wallis and Dunn's multiple comparison test (K).
